# Development and validation of a phenotypic high-content imaging assay for assessing the antiviral activity of small-molecule inhibitors targeting the Zika virus

**DOI:** 10.1101/302927

**Authors:** Jean A. Bernatchez, Zunhua Yang, Michael Coste, Jerry Li, Sungjun Beck, Yan Liu, Alex E. Clark, Zhe Zhu, Lucas A. Luna, Christal D. Sohl, Byron W. Purse, Rongshi Li, Jair L. de Siqueira-Neto

## Abstract

Zika virus (ZIKV) has been linked to the development of microcephaly in newborns, as well as Guillain-Barré syndrome. There are currently no drugs available to treat infection, and accordingly there is an unmet medical need for discovery of new therapies. High-throughput drug screening efforts focusing on indirect readouts of cell viability are prone to a higher frequency of false positives in cases where the virus is viable in the cell but the cytopathic effect is reduced or delayed. Here, we describe a fast and label-free phenotypic high-content imaging assay used to detect cells affected by the viral-induced cytopathic effect (CPE) using automated imaging and analysis. Protection from CPE correlates with a decrease in viral antigen production as observed by immunofluorescence. We trained our assay using a collection of nucleoside analogues against ZIKV; the previously reported antiviral activities of 2’-C-methylribonucleosides and ribavirin against the Zika virus in Vero cells were confirmed using our developed method. Profiling of a novel library of 24 natural product derivatives using our assay revealed compound **1** as an inhibitor of ZIKV-induced cytopathic effect; activity of the compound was confirmed in human fetal neural stem cells (NSCs). The described technique can be easily leveraged as a primary screening assay for profiling large compound libraries against ZIKV, and can be expanded to other ZIKV strains and other cell lines displaying morphological changes upon ZIKV infection.

## Introduction

The reemergence of the Zika virus (ZIKV) in recent years as an infectious agent of global concern has driven extensive scientific investigation into the pathology and treatment of infection (1, 2). This pathogen from the viral family *Flaviviridae* is transmitted by *Aedes* sp. mosquitos and is associated with microcephaly in newborns and neural-inflammatory diseases such as Guillain-Barré syndrome and ophthalmological complications in adults (3, 4). No approved therapy is currently available to treat this infection, stressing the importance of developing new vaccines and antivirals.

Viral polymerases remain attractive targets in the development of antivirals due to their proven clinical usefulness and their essential activity in viral life cycles (5–7). Nucleoside analogues abrogate nucleic acid synthesis and consequently viral genome replication when incorporated by the viral polymerase and represent an important component of treatment for numerous viral infections. Drug repurposing studies have recently identified numerous candidate inhibitors of ZIKV replication (8, 9), including the nucleoside analogue ProTide (prodrug nucleotide) sofosbuvir, a direct-acting antiviral against the Hepatitis C virus (HCV) (10, 11). Other nucleoside analogues that have been investigated for anti-ZIKV activity (targeting the viral polymerase or nucleoside biosynthesis) include 7-deaza-2´-C-methyladenosine (7-deaza-2’-CMA) (12, 13), the adenosine analogue NITD008 (14), 2’-C-methylribonucleosides (13, 15, 16), 3’-O-methylribonucleosides (13), ribavirin (17) and 5-fluorouracil (18).

Testing antiviral activity in cell-based phenotypic assays has been conducted in many of these studies. Readouts of antiviral activity used to assess compound efficacy and toxicity include qRT-PCR of viral RNA (19), plaque reduction assays (13), cell-viability assays (18), caspase-activation assays (20), luciferase Zika virus (21), Zika replicons (22) and immunofluorescence-based detection of the virus (9). While these assays are robust, most methods are either too labor-intensive or prohibitively expensive for primary large-scale compound library screening. Furthermore, protection from viral-induced cell morphology changes, or cytopathic effects (CPE) (23), is inferred from indirect measurements. Morphological confirmation of antiviral affect in cellular systems has become increasingly used in drug development in recent years due to technological innovations in robotics, imaging and automated image analysis, as these screens are target agnostic and can help discover “first-in-class” inhibitors (24). In this work, we developed a cell-based phenotypic assay using automated image segmentation and analysis to assess the CPE of the recent ZIKV clinical isolate from Panama H/PAN/2016/BEI-259634, NR-50210 (GenBank: <u>KX198135</u>) in Vero cells. We then trained our developed method for detecting antiviral activity using a library of nucleoside analogues. Subsequently, we tested the method using a novel set of marine natural product derivatives and discovered a new chemical scaffold with anti-Zika inhibitory activity.

## Results and Discussion

### Assessment of CPE

Cells were cultured in the presence and absence of ZIKV in 1536-well plates for 72 hours at multiplicity of infection (MOI) of 10 to ensure sufficient numbers of detached cells from the monolayer on the plate surface during viral infection. Bright-field images of the wells were acquired using an ImageXpress Micro high-content imager with 10x objective lens (Molecular Devices) (Fig. 1a). Cells were subsequently fixed with 4% formaldehyde and stained with 4’,6-diamidino-2-phenylindole (DAPI). The plates were re-imaged to acquire pictures of the cell nuclei (Fig. 1b). Images were then analyzed using a custom module in MetaXpress (Molecular Devices) to generate segmentation counts of detached cells in bright-field, as well as segmentation counts of total cell nuclei (Fig. 1c and d). Percent levels of CPE were determined by dividing the number of detached cells by the total number of cell nuclei and multiplying by 100. Percent CPE was normalized to the uninfected controls by subtracting the percent CPE in uninfected controls from infected wells. Using 128 positive and 128 negative control wells, a Z’ value of 0.50 was obtained, which is comparable to those reported for other phenotypic assays used to screen for ZIKV inhibitors (8, 9, 18), and indicates that the assay is useful as a primary screening tool for large compound libraries. Our method requires only one short staining step (DAPI) and two sets of images to be taken to assess antiviral activity, reducing the amount of sample manipulation and allowing for same-day data acquisition.

**Figure 1.**
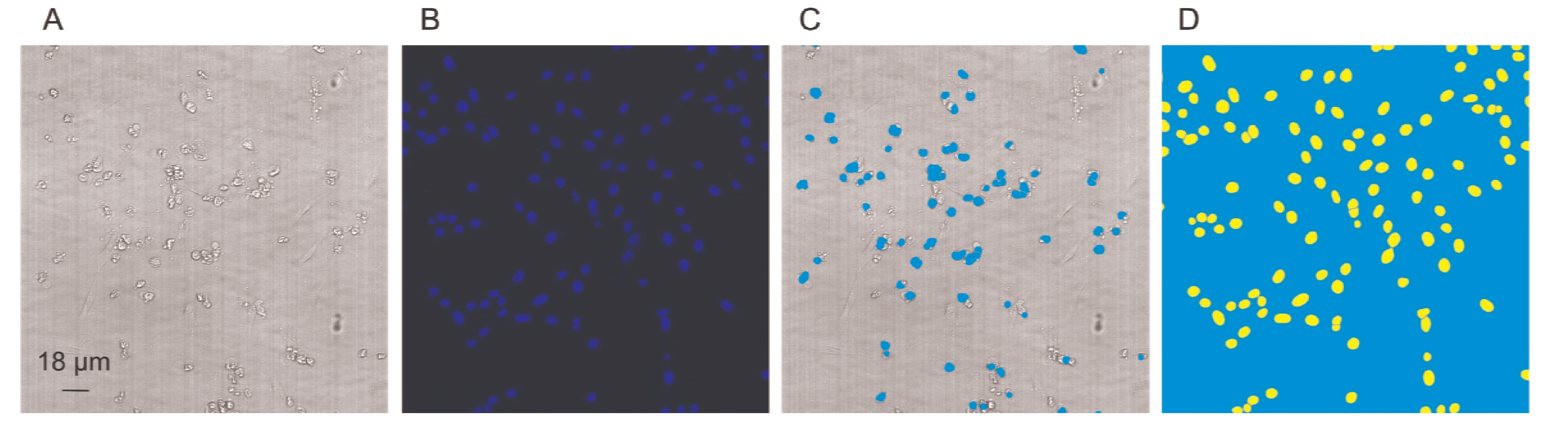
Automated segmentation analysis of Vero cells undergoing ZIKV-induced CPE and total cell nuclei. Detached cells and total cell nuclei were acquired at 10x magnification and tabulated using an automated image analysis protocol. A) Bright-field image of cells infected with MOI 10 H/PAN/2016/BEI-259634 ZIKV 72 hours post-infection. B) Segmentation of detached cells using an automated image analysis module. C) Image of DAPI-stained Vero cells. D) Segmentation of cell nuclei using an automated image analysis module.

### Training of antiviral assay using published nucleoside analogues

We assessed the antiviral activity of a small training library of nucleoside analogues and nucleoside analogue ProTides (25) (Fig. 2). Our selection of nucleoside analogues for testing encompasses three types of structural features known to inhibit viral polymerases by alternative mechanisms. The first category is obligate chain terminators, which lack the 3’ hydroxyl group needed for elongation after incorporation into a nascent RNA strand (13, 15). We selected compounds that lack the 3’ hydroxyl (i.e. replacing it with hydrogen) or that substitute a methoxy or fluoro group in its place (Table 1 and Fig. 2). While having the potential for high potency against enzymatic RNA synthesis when delivered as a nucleoside triphosphate, the 3’ modifications can attenuate nucleoside phosphorylation in cells (26). The second category is non-obligate chain terminators, which typically include 2’ modifications that block chain elongation for conformational or steric reasons (11, 27). In this category, we included the 2’-C-methyl compounds previously shown to be active against ZIKV in cell-based assays, plus additional modifications. Last, we included nucleobase modifications associated with antiviral activity by lethal mutagenesis (28, 29). While typically potent, these inhibition mechanisms are often associated with greater cytotoxicity. Since some of the structural modifications in these substrate analogues are known to be associated with reduced nucleo(s/t)ide kinase compatibility, we prepared ProTide prodrugs for a selection of the compounds (25). The ProTide nucleoside monophosphate masking strategy is used in the clinically approved anti-HCV drug sofosbuvir (11).

**Figure 2.**
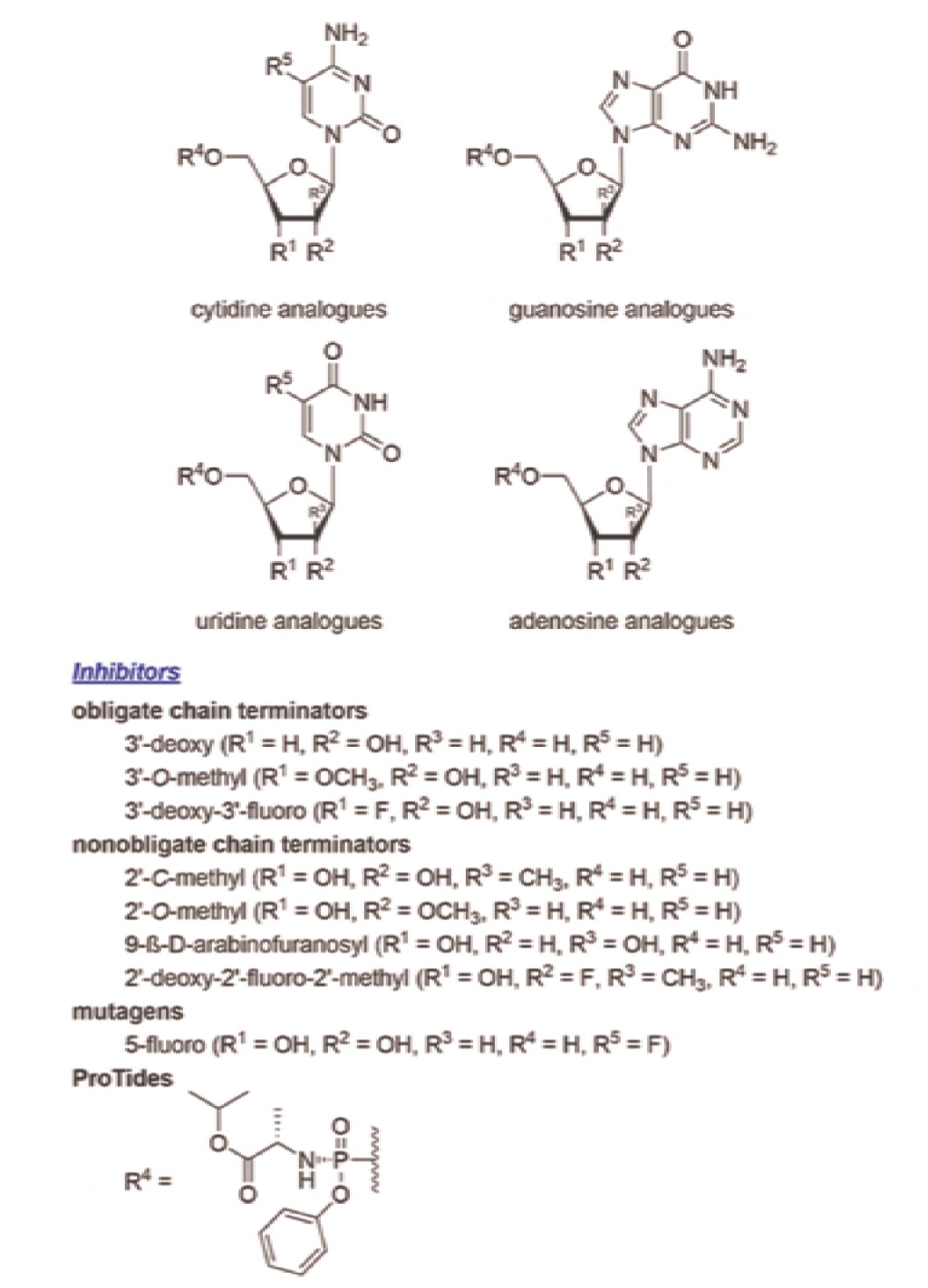
Structural modifications of polymerase substrate analogue inhibitors tested against Zika virus in this phenotypic high-content imaging assay.

**Table 1.** EC_50_ and CC_50_ for a selected library of nucleoside analogues using the developed phenotypic assay. Values are the result of two independent biological replicates with (± Standard Error). The selectivity index is calculated by dividing the CC50 by the EC50.

| Compound Name | EC<sub>50</sub> (μM) | CC<sub>50</sub> (μM) | Selectivity Index (SI) |
| --- | --- | --- | --- |
| 2'-C-Methylcytidine | 0.30 (±0.12) | >100.00 | >300 |
| 2'-C-Methyladenosine | 0.60 (±0.25) | >100.00 | >150 |
| 2'-C-Methylguanosine | 2.20 (±0.49) | >100.00 | >40 |
| 2'-C-Methyluridine | 4.21 (±0.96) | >100.00 | >20 |
| Ribavirin | 20.80 (±4.45) | >100.00 | >4 |
| 5-Fluoro-2'-deoxyuridine | <0.20 | <0.20 | - |
| 5-Fluorouridine | 1.11 (±0.33) | <0.20 | <0.2 |
| 5-Fluorocytidine | 1.92 (±0.82) | <0.20 | <0.1 |
| Sofosbuvir | >100.00 | >100.00 | - |
| 2'-O-Methylcytidine | >100.00 | >100.00 | - |
| 3'-Deoxyadenosine | >100.00 | >100.00 | - |
| 3'-Deoxycytidine | >100.00 | 79.60 (±37.57) | <0.8 |
| 3'-O-Methylguanosine | >100.00 | >100.00 | - |
| 3'-O-Methyluridine | >100.00 | >100.00 | - |
| 9-(β-D-Arabinofuranosyl)guanine | >100.00 | >100.00 | - |
| 2'-C-Methylcytidine ProTide | >100.00 | >100.00 | - |
| 2'-C-Methyluridine ProTide | >100.00 | >100.00 | - |
| 2'-C-Methyladenosine ProTide | >100.00 | >100.00 | - |
| 3'-Deoxyadenosine ProTide | >100.00 | >100.00 | - |
| 5'-Fluorocytidine ProTide | >100.00 | >100.00 | - |
| 3'-O-Methyluridine ProTide | >100.00 | >100.00 | - |
| 2'-O-Methylcytidine ProTide | >100.00 | >100.00 | - |
| 3'-Deoxycytidine ProTide | >100.00 | >100.00 | - |
| 3'-Deoxy-3'-fluoroguanosine<br>ProTide | >100.00 | >100.00 | - |

Compounds were pre-spotted onto plates using an Acoustic Transfer System instrument (EDC Biosciences) in a ten point, two-fold dose response from 100 μM to 0. 197 μM. Vero cells and ZIKV were then added to the plate as described above and incubated with compound for 72 hours. Bright-field and DAPI channel images were acquired as described above; normalized percent activity was plotted versus concentration of compound and fit to an EC_50_ function (CDD Vault) (Figure 3). Percent cell viability upon compound treatment was determined by dividing the number of cell nuclei in a compound-treated well by the average number of cell nuclei in control wells treated with vehicle (0.1% dimethyl sulfoxide (DMSO)) and multiplying by 100. The obtained percent cell viability values were then plotted against compound concentration and fit to a CC_50_ function (CDD Vault). The results obtained from our assay demonstrate that the previously studied ribavirin (EC_50_ = 20.8 μM) and 2’-C-methylribonucleosides, with 2’-C-methylcytidine being the most potent (EC_50_= 0.297 μM), are inhibitors of ZIKV replication (Table 1). No cell toxicity was observed for these compounds up to concentrations of 100 μM. None of the other nucleoside analogues tested displayed specific antiviral activity in our assay, and 5-fluorouracil, 5-fluorocytidine and 5-fluoro-2’– deoxyuridine displayed high levels of cellular toxicity (CC_50_ < 0.197 μM).

**Figure 3.**
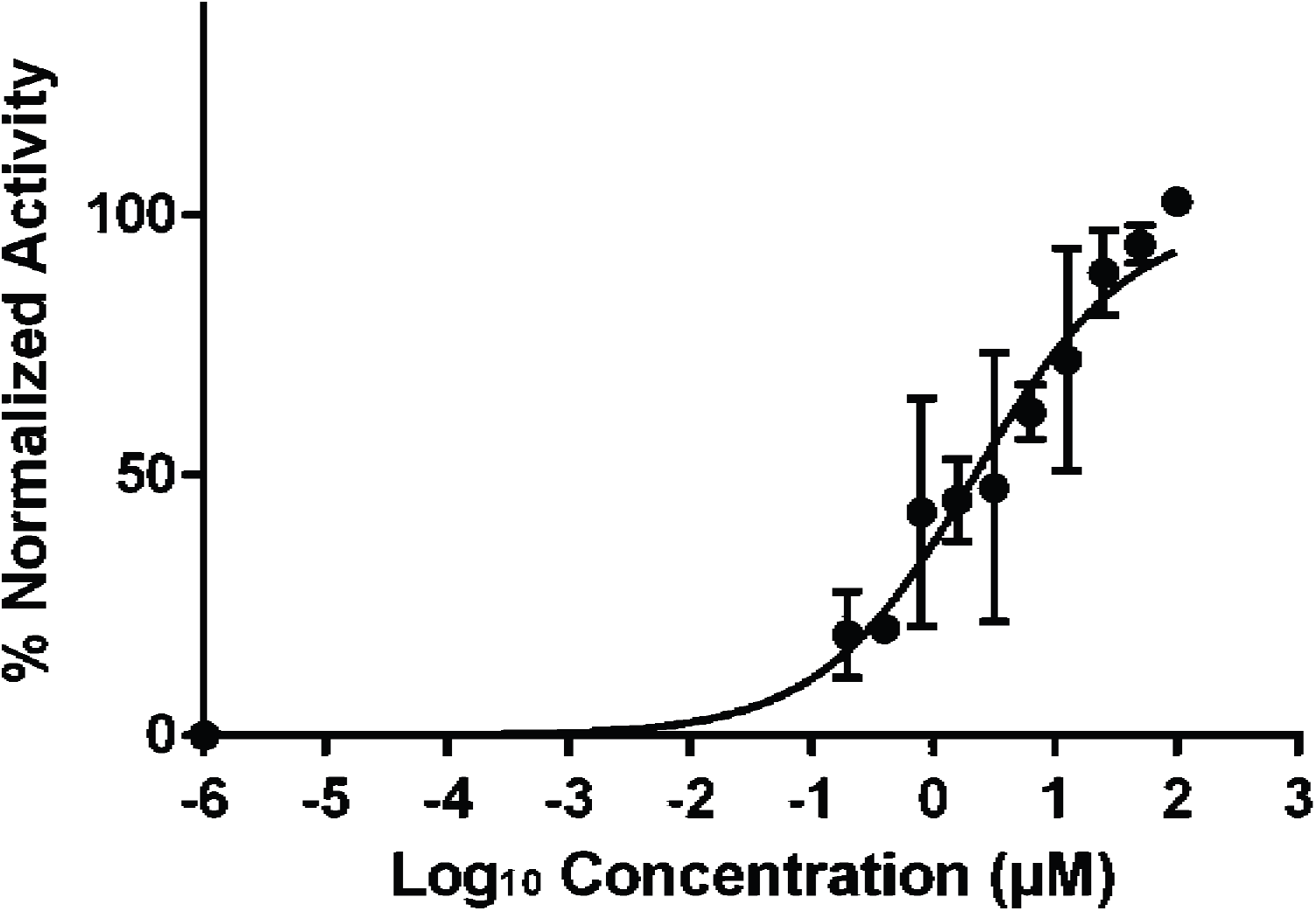
Sample EC_50_ curve of inhibitor dose-response data acquired from the phenotypic screen. Concentration of inhibitor was plotted against normalized activity (Norm Activity (%)) and an EC_50_ curve fit was applied to the dose-response data for the ZIKV inhibitor 2’-C-methylguanosine (CDD Vault and GraphPad Prism 6). The experiment was repeated in duplicate using two different passages of Vero cells.

### Secondary assay counter-screen

We counter-screened our remaining hit compounds by staining the plates for ZIKV antigen via immunofluorescence (Fig. 4) to confirm reduction in virus production upon compound treatment. Reduction in the percentage of detached cells observed in bright-field upon treatment with the potent 2’-C-methylcytidine correlated with a reduction in the amount of viral antigen, indicating agreement between our image-based methods and previously published assays.

**Figure 4.**
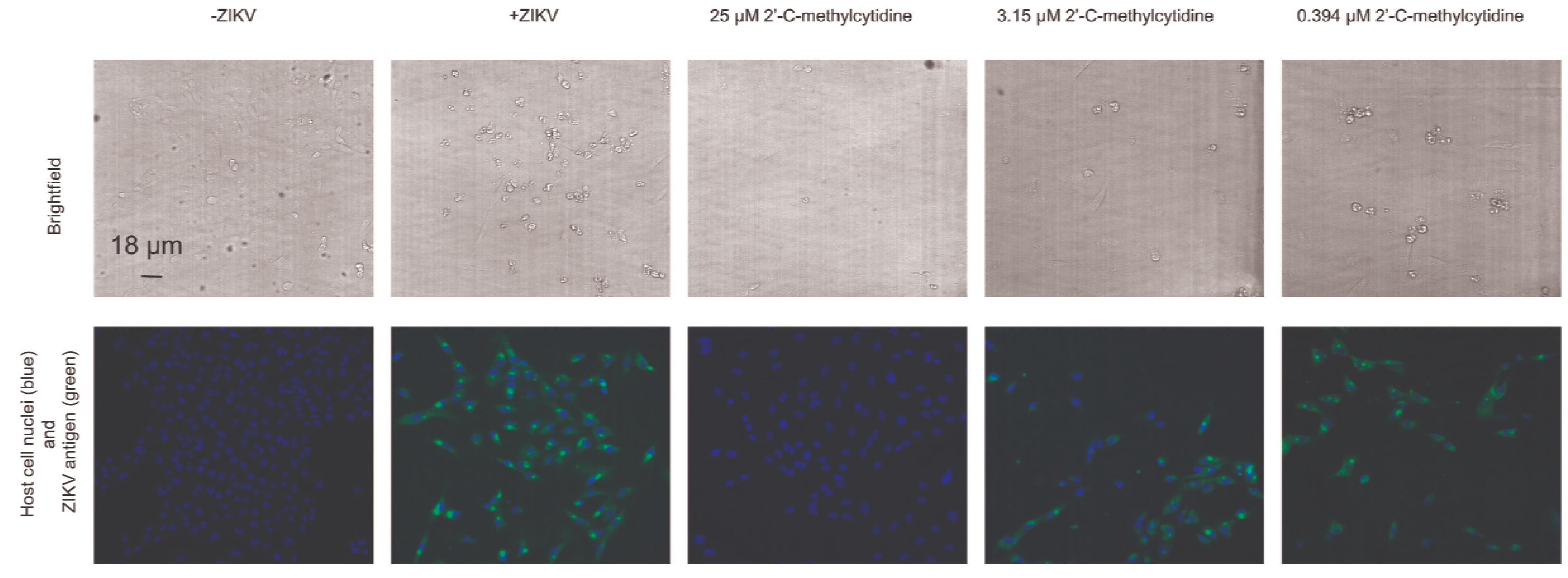
2’-C-methylcytidine inhibits cellular replication of ZIKV and blocks the cytopathic effect of the virus in Vero cells. 2’-C-methylcytidine was tested in our phenotypic CPE assay and counter-screened using immunofluorescence. CPE-affected cells (bright-field), cell nuclei (blue) and ZIKV antigen (green) image acquisition was performed at 10x magnification for uninfected cells, cells infected with MOI 10 H/PAN/2016/BEI-259634 ZIKV for 72 hours in presence of vehicle (DMSO), and cells infected with MOI 10 H/PAN/2016/BEI-259634 ZIKV for 72 hours in the presence of varying concentrations of 2’-C-methylcytidine.

In contrast with previously published reports that ribavirin has no effect on preventing CPE in Vero cells (8), we observed a reduction in the number of detached cells during ZIKV infection, as well as a reduction in ZIKV antigen production via immunofluorescence when cells were treated with ribavirin at high concentrations (Fig. 5). This is most likely due to our direct visualization of CPE through cell imaging techniques and the higher test concentration of ribavirin (up to 100 μM) used in this study.

**Figure 5.**
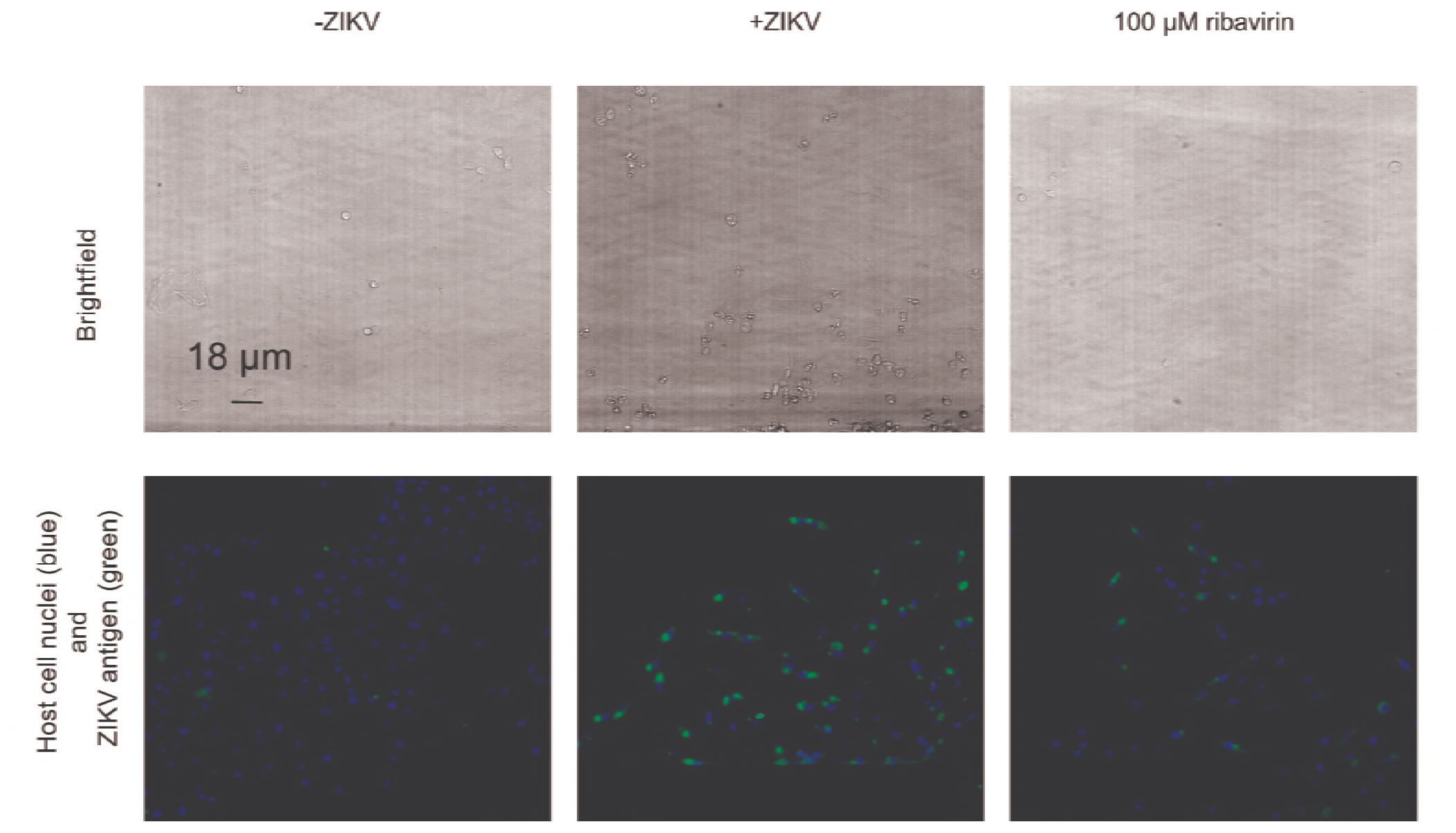
Ribavirin reduces production of ZIKV antigen and blocks the cytopathic effect of the virus in Vero cells. Shown are the results for the testing of ribavirin in our phenotypic CPE assay and counter-screening of the compound using immunofluorescence. CPE-affected cells (bright-field), cell nuclei (blue) and ZIKV antigen (green) image acquisition was performed at 10x magnification for uninfected cells, cells infected with MOI 10 H/PAN/2016/BEI-259634 ZIKV for 72 hours in presence of vehicle (DMSO), and cells infected with MOI 10 H/PAN/2016/BEI-259634 ZIKV for 72 hours in the presence of 100 μM ribavirin.

Interestingly, we found that sofosbuvir, which has been reported to inhibit ZIKV (10, 11, 30), as well as ProTides of 2’-C-methylribonucleosides, were inactive up to concentrations of 100 μM. Recent studies have demonstrated that sofosbuvir (30) and ribavirin (19) have cell line-dependent inhibitory profiles. Our results corroborate the finding that sofosbuvir is inactive in the Vero cell line (30). It is conceivable that, like sofosbuvir, poor metabolic activation of the 2’-C-methylribonucleoside ProTides in Vero cells results in little activity. We are currently developing image analysis modules to expand our phenotypic assay to cell lines of neural lineage to assess antiviral activity of these compounds in different metabolic environments.

### Screening of a novel library of natural products-derivatives using the developed assay

Although the design and synthesis will be reported elsewhere, here we briefly describe our fragment-based approach towards the natural product-based small-molecule library used to validate our screening assay. We recently designed, synthesized and reported marinopyrrole derivatives as potent anti-methicillin resistant Staphylococcus aureus (MRSA) and anticancer agents (31–38). As shown in Figure 6, marinopyrroles 1a and 1b have clogP values of 6.5 and 6.1, respectively, which violates Lipinski’s rule of five (39). In order to improve physicochemical properties, we designed a library containing 24 members with lower clogP values ranging from 2.1 to 5.0. We performed structural simplification and optimization of these marinopyrrole derivatives and came up with a novel series of pyrrolomycin-like derivatives (40, 41). Compound **1** (Fig. 6), as one of the library members with a clogP value of 4.9, was fully characterized using NMR and high resolution mass spectrometry.

**Figure 6.**
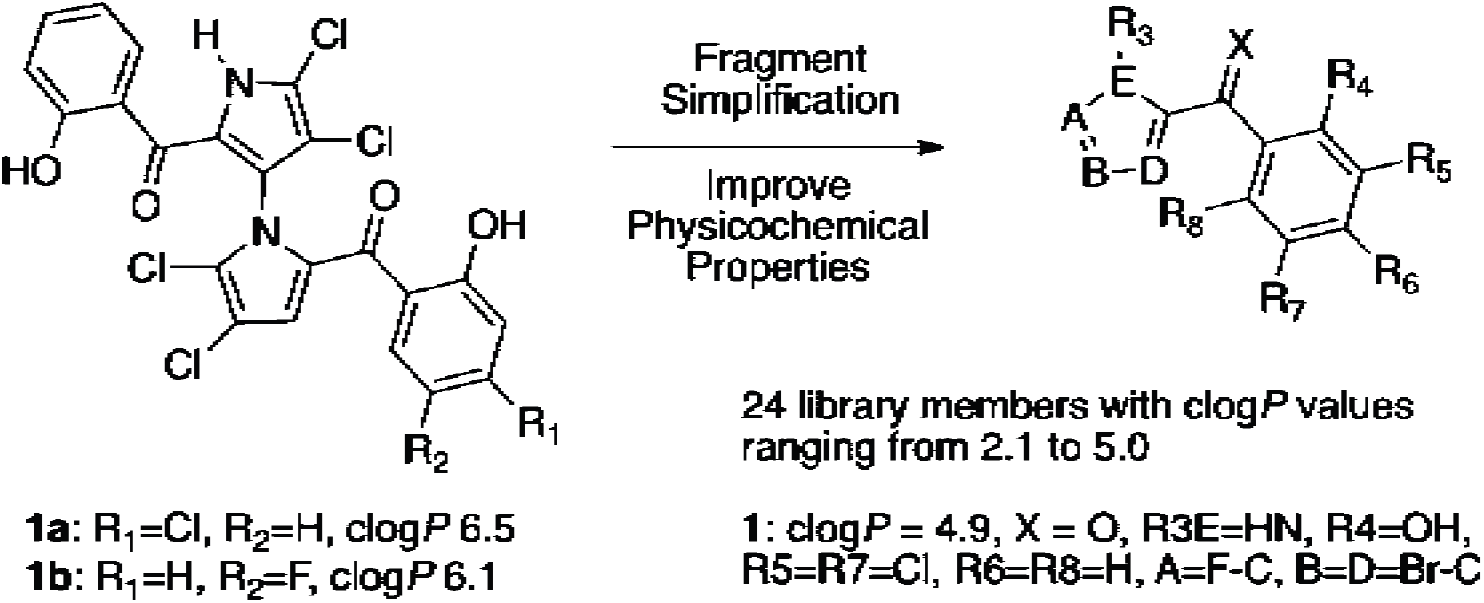
A library of natural product derivatives resulted from structural optimization of marinopyrroles.

Validation of the assay as a platform for the identification of new chemical entities with anti-ZIKV activity was conducted using the library of 24 natural products (Fig. 6). Screening of the library revealed compound **1** as being able to prevent virus-induced CPE, with an EC_50_ of 5.95 μM and CC_50_ of > 10 μM (Table 2); all 23 other compounds in the library were inactive in our primary screening assay up to concentrations of 10 μM. Reduction in viral antigen production upon treatment with compound **1** was also observed in the immunofluorescence assay (Fig. 7). Lastly, we tested compound **1** in a luminescence-based cell survival assay (Promega’s Cell Titer Glo) in a human fetal neural stem cell (NSC) model of ZIKV infection. The compound was able to protect cells from virus-induced cell death, with an EC_50_ of 8.56 μM and CC_50_ of > 10 μM (Table 2).

**Figure 7.**
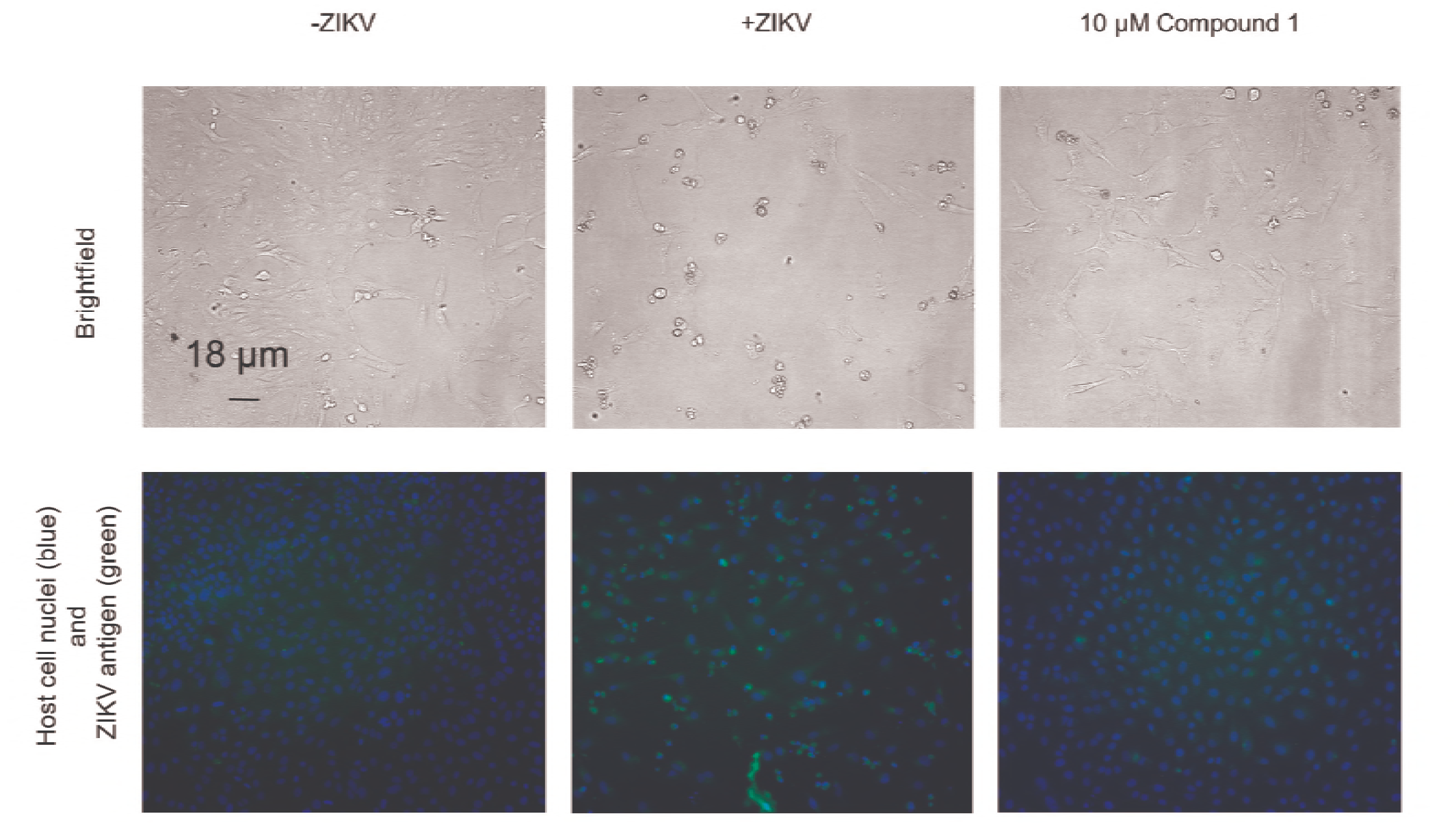
Compound **1** has anti-ZIKV activity in Vero cells. Compound **1** was assessed in the phenotypic CPE assay and counter-screened using immunofluorescence. Images were acquired for CPE-affected cells (brightfield), cell nuclei (blue) and ZIKV antigen (green) image acquisition at 10x magnification for uninfected cells, cells infected with MOI 10 H/PAN/2016/BEI-259634 ZIKV for 72 hours in presence of vehicle (DMSO), and cells infected with MOI 10 H/PAN/2016/BEI-259634 ZIKV for 72 hours in the presence of 10 μM compound 1.

**Table 2.** EC_50_ and CC_50_ for compound **1** using the developed phenotypic assay and Cell Titer Glo assay. Values are the result of two independent biological replicates with (± Standard Error). ^a^Phenotypic assay. ^b^Cell Titer Glo assay.

| Compound<br>Name | EC<sub>50</sub> (μM)<br>PA<sup>a</sup> | CC<sub>50</sub> (μM)<br>PA | Selectivity<br>Index (SI)<br>PA | EC<sub>50</sub> (μM)<br>CTG<sup>b</sup> | CC<sub>50</sub> (μM)<br>CTG | Selectivity<br>Index (SI)<br>CTG |
| --- | --- | --- | --- | --- | --- | --- |
| Compound 1 | 5.95<br>(±0.72) | >10.00 | >1.6 | 8.56<br>(±0.70) | >10.00 | >1.1 |

### Conclusion

We have developed a simple and rapid phenotypic screen that can be used as a primary screening tool for the profiling of large compound libraries against ZIKV. This method can be expanded to other cell lines which display morphological changes during ZIKV infection that can be detected by automated image analysis software. As proof-of-concept, we confirmed that 2’-C-methylribonucleosides and ribavirin can protect Vero cells from ZIKV-induced cytopathic effect, and identified compound **1** as a novel scaffold against ZIKV using our assay. Future studies will focus on structural-activity-relationship (SAR) optimization of compound **1** for further development and screening large-scale libraries to identify additional novel chemical entities for the inhibition of ZIKV replication.

## Materials and Methods

### Nucleoside chemistry

All reagents and chemicals used were purchased from Acros Organics, and Fisher Chemical at ACS grade or higher quality and used as received without further purification. All nucleoside analogues were obtained from Carbosynth LLC and N-[(S)-(2,3,4,5,6-pentafluorophenoxy)phenoxyphosphinyl]-L-alanine 1-Methylethyl ester was obtained from AK Scientific. All reactions were carried out in oven-dried Schlenk tubes under nitrogen atmosphere, using commercially available anhydrous solvents and monitored by thin layer chromatography, detected by UV light. ^1^H NMR spectra were acquired on a Varian 400 MHz and a Varian 500 MHz NMR spectrometer and recorded at 298 K. Chemical shifts were referenced to the residual protio solvent peak and given in parts per million (ppm). Splitting patterns are denoted as s (singlet), d (doublet), dd (doublet of doublet), ddd (doublet of doublet of doublet), t (triplet), q (quartet), m (multiplet).

### Synthesis of nucleoside analogue ProTides

To a stirred suspension of nucleotide (0.09 mmol, dried under vacuum at 50 °C overnight) in dry THF (1 mL) was added a 2.0 M solution of *isopropyl magnesium* chloride in THF (96 μL, 0.19 mmol). The mixture was stirred at 0 °C for 30 min, then allowed to warm to room temperature and stirred for an additional 30 min. The reaction mixture was then cooled to 0 °C and N-[(S)-(2,3,4,5,6-pentafluorophenoxy)phenoxyphosphinyl]-L-alanine 1-methylethyl ester (46 mg, 0.10 mmol) was added. The reaction mixture was stirred for 18 h as the temperature was allowed to warm to room temperature. The solvent was removed by rotary evaporation. The reaction mixture was purified using first flash chromatography (0 to 30% MeOH in dichloromethane gradient) and then preparative, normal-phase HPLC (10 to 40% MeOH in dichloromethane gradient) to afford nucleoside analogue ProTide as assessed by NMR.

### Cells

Vero cells were purchased from ATCC (ATCC CCL-81) and cultured in Dulbecco’s Modified Eagle Medium (DMEM) (Gibco) supplemented with 10% Fetal Bovine Serum (FBS) (Sigma), 4.5 g/L D-Glucose, 4 mM L-Glutamine and 110 mg/L sodium pyruvate (Gibco). Culturing of cells was conducted in incubators at 37 C and 5% CO_2_. Human fetal NSCs were obtained commercially from Clontech (Human Neural Cortex (Y40050). The NSCs were maintained in Neurobasal®-A without phenol red (Thermo Fisher) with the addition of B27 supplement (1:100, Thermo Fisher, #12587010), N2 supplement (1:200, Invitrogen, #17502-048), 20 ng/ml FGF (R&DSystems 4114-TC-01M), 20 ng/ml EGF (R&D Systems 236-EG-01M), GlutaMax (Thermo Fisher, #35050061), and sodium pyruvate. The cells were plated into 6 well plates pre-coated with laminin (10 mg/ml, Sigma #-L2020). Cells were grown to near confluency (80-90%) prior to passage. For passaging, cells were rinsed gently with PBS (without calcium and magnesium) and then accutase (Sigma, A6964) was added for 5 min at 37 C to allow detachment.

### Viruses

The following material was obtained through BEI Resources, NIAID, NIH: Zika Virus, H/PAN/2016/BEI-259634, NR-50210 (GenBank: KX198135). The virus was expanded in Vero cells for 2-3 serial passages to amplify titers. Infected cell supernatants were centrifuged to remove cell debris and then concentrated through a sucrose cushion. Concentrated virus was resuspended in neural maintenance medium base (50% DMEM/F12-Glutamax, 50% Neurobasal medium, 1x N-2 Supplement, 1x B-27 Supplement (Life Technologies)) supplemented with 1% DMSO (Sigma) and 5% FBS (Gibco) and stored at −80 ºC. Viral stock titers were determined by plaque assay on Vero cells and were greater or equal to 2 × 10^8^ plaque forming units (pfu)/mL.

### Cytopathic Effect Assay

Compounds were pre-spotted to 1536 Greiner black well plates using an Acoustic Transfer System (EDC Biosciences). Vero cells were seeded at 100 per well in the presence of 0.1% DMSO or test compound, and with ZIKV at MOI 10 or without virus. Cultures were incubated for 72 hours in incubators at 37 ºC and with 5% CO_2_. Plate wells were then imaged in bright-field using an ImageXpress microplate imager (Molecular Devices), cells were subsequently fixed with 4% formaldehyde, stained with DAPI in 0.85% NaCl, 0.1% Triton-X, 0.01% sodium azide and 100 mM ammonium chloride for **1** hour, and then imaged to acquire images of cell nuclei. A custom analysis module in MetaXpress software (Molecular Devices) was used to segment and tabulate total numbers of detached cells in bright-field and total cell nuclei (see below). Data processing, including normalization, EC50 and CC50 calculations, was conducted using CDD Vault – Collaborative Drug Discovery Inc. (Burlingame, CA) and GraphPad Prism (La Jolla, CA).

### Automated Image Analysis Module

Using the MetaXpress Custom Module Editor, an image segmentation protocol was created to quantify Vero cells undergoing CPE during ZIKV infection in bright-field: a simple threshold from 0 to 65535 intensity units was created. Next, a Gaussian filter with a sigma value of 2 was first applied to the image. Using the multiply function, the product of these two images was generated and then multiplied by 5000. A gradient was then applied to this result using a pixel size of 2. An Open filter was then applied with a circle filter shape and pixel size of 2, with greyscale reconstruction included. Subsequently, a Find Round Objects filter was used to detect detached Vero cells in the image; approximate object minimum was set to 10 μm, approximate object maximum was set to 30 μm, and intensity above local background was 300 intensity units. A Filter Mask was then created, which was used in the final object quantification: the breadth of the detected round objects was subjected to a RangeFilter function of 10 to 30 μm, inclusive, while average object intensity was treated to a RangeFilter function of 2000 to 60000 intensity units.

To normalize the number of objects detected in bright-field to the number of cells in a given image, a cell nuclei detection mask for DAPI-stained cells was developed using the MetaXpress Custom Module Editor. An Open filter for circular objects with size of 9 pixels was first applied to the images of DAPI-stained cells. A Gaussian filter of sigma value 2 was then applied to the resultant image. A Find Round Objects filter was next used to detect nuclei, with approximate object minimum set to 6 μm, approximate object maximum set to 25 μm, and intensity above local background of 30 intensity units.

### Immunofluorescence

Following bright-field and DAPI imaging, plates were incubated with anti-flavivirus group antigen primary antibody, clone D1-4G2-4-15 (Millipore) at 1:200 stock dilution overnight at 4 ºC. Upon aspiration of primary antibody from the plates, FITC goat anti-mouse IgG (H+L) secondary antibody (Life Technologies) was added at 1:1000 stock dilution and incubated for **1** hour at 25 ºC. Secondary antibody was aspirated; samples were then incubated in a buffer containing 0.85% NaCl, 0.1% Triton-X, 0.01% sodium azide and 100 mM ammonium chloride and imaged in FITC channel using an ImageXpress Micro plate imager (Molecular Devices).

### Cell Titer Glo assay

NSCs (5000 cells/well) were seeded into 384-well plates 12 h prior to infection. ZIKV HPAN was added at MOI 5 (5 FFU/cell). In all experiments, mock infected cells were incubated in parallel. Cell viability was measured using Cell Titer-Glo (Promega) and readouts for luminescence intensity were conducted using an EnVision plate reader (Perkin Elmer). All data were normalized to DMSO controls and expressed as a relative luminescence intensity.

## Acknowledgments

This research did not receive any specific grant from funding agencies in the public, commercial, or not-for-profit sectors. B.W.P, M.C., L.A.L., and C.D.S. thank San Diego State University for financial support. Design and synthesis of a library of natural product derivatives were partially supported by Nebraska Research Initiative (NRI) and Start-up funds to R.L. by University of Nebraska Medical Center. Screening experiments were developed at the UCSD Screening Core.

